# The genotype-phenotype landscape of an allosteric protein

**DOI:** 10.1101/2020.07.10.197574

**Authors:** Drew S. Tack, Peter D. Tonner, Abe Pressman, Nathanael D. Olson, Sasha F. Levy, Eugenia F. Romantseva, Nina Alperovich, Olga Vasilyeva, David Ross

**Affiliations:** National Institute of Standards and Technology, Gaithersburg, MD, 20899, USA; SLAC National Accelerator Laboratory, Menlo Park, CA, 94025, USA; Joint Initiative for Metrology in Biology, Stanford, CA, 94305, USA

## Abstract

Allostery is a fundamental biophysical mechanism that underlies cellular sensing, signaling, and metabolism. Quantitative methods to characterize the genotype-phenotype relationships for allosteric proteins would provide data needed to improve engineering of biological systems, to uncover the role of allosteric mis-regulation in disease, and to develop allosterically targeted drugs^1^. Here we report the large-scale measurement of the genotype-phenotype landscape for an allosteric protein: the *lac* repressor from *Escherichia coli*, LacI. Using a method that combines long-read and short-read DNA sequencing, we quantitatively determine the dose-response curves for nearly 10^5^ variants of the LacI sensor. With the resulting data, we train a deep neural network (DNN) capable of predicting the dose-response curves for additional LacI genotypes *in silico.* We then map the impact of amino acid substitutions on the allosteric function of LacI. Additionally, we demonstrate engineering of allosteric function with unprecedented accuracy by identifying LacI variants from the measured landscape with quantitatively specified dose-response curves. Finally, we discover sensors with previously unreported band-stop dose-response curves. Overall, our results provide the first high-coverage, quantitative view of genotype-phenotype relationships for an allosteric protein, revealing a surprising diversity of phenotypes and showing that each phenotype is accessible via multiple distinct genotypes.

## Main

Genetic sensors are allosteric proteins that regulate gene expression in response to stimuli, giving cells the ability to regulate their metabolism and respond to environmental changes. Genetic sensors are also central to engineering biology, with applications in programming logic^2^, optimizing biosynthetic pathways^3,4^, and controlling cellular differentiation^5^. Like many allosteric genetic sensors, LacI binds to DNA upstream of regulated genes, preventing transcription. LacI regulates gene expression by switching between DNA-binding and non-binding conformations upon ligand binding at an allosteric site. This allosteric switching is determined by several biophysical constants, including ligand-binding affinity, DNA-binding affinity, and the allosteric constant (the thermodynamic equilibrium between the two conformations)^6–8^. These constants depend on amino acid residues and interactions spread widely across the protein, making it difficult to predict the effects of changes to the protein sequence.

Advances in DNA sequencing have enabled the phenotypic characterization of 10^4^ to 10^5^ genotypes in a single measurement^9–14^. The resulting large-scale genotype-phenotype landscapes have increased our understanding of biological function and evolutionary dynamics and improved our ability to engineer biology. Notably, measurements at this scale facilitate the exploration of genotypes with mutations widely spread throughout a nucleotide or amino acid sequence. So, genotype-phenotype landscape measurements are ideal for probing complex biological mechanisms, like allostery, that emerge from broadly distributed intramolecular interactions.

### Mapping the genotype-phenotype landscape

Dose-response curves describe the phenotype of a genetic sensor by relating the concentration of an input ligand (*L*) to an output response (gene expression, *G*). Genetic sensors typically have sigmoidal dose-response curves that can be represented with the Hill equation:

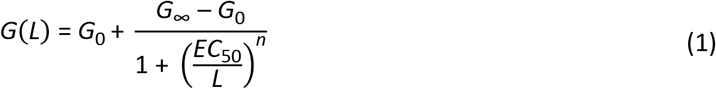

where *G*_0_ is gene expression in the absence of ligand, *G*_∞_ is gene expression at saturating ligand concentrations, *EC*_50_ is the ligand concentration that gives half-maximal response, and *n* quantifies the steepness of the dose-response curve (Fig. 1).

**Fig. 1.**
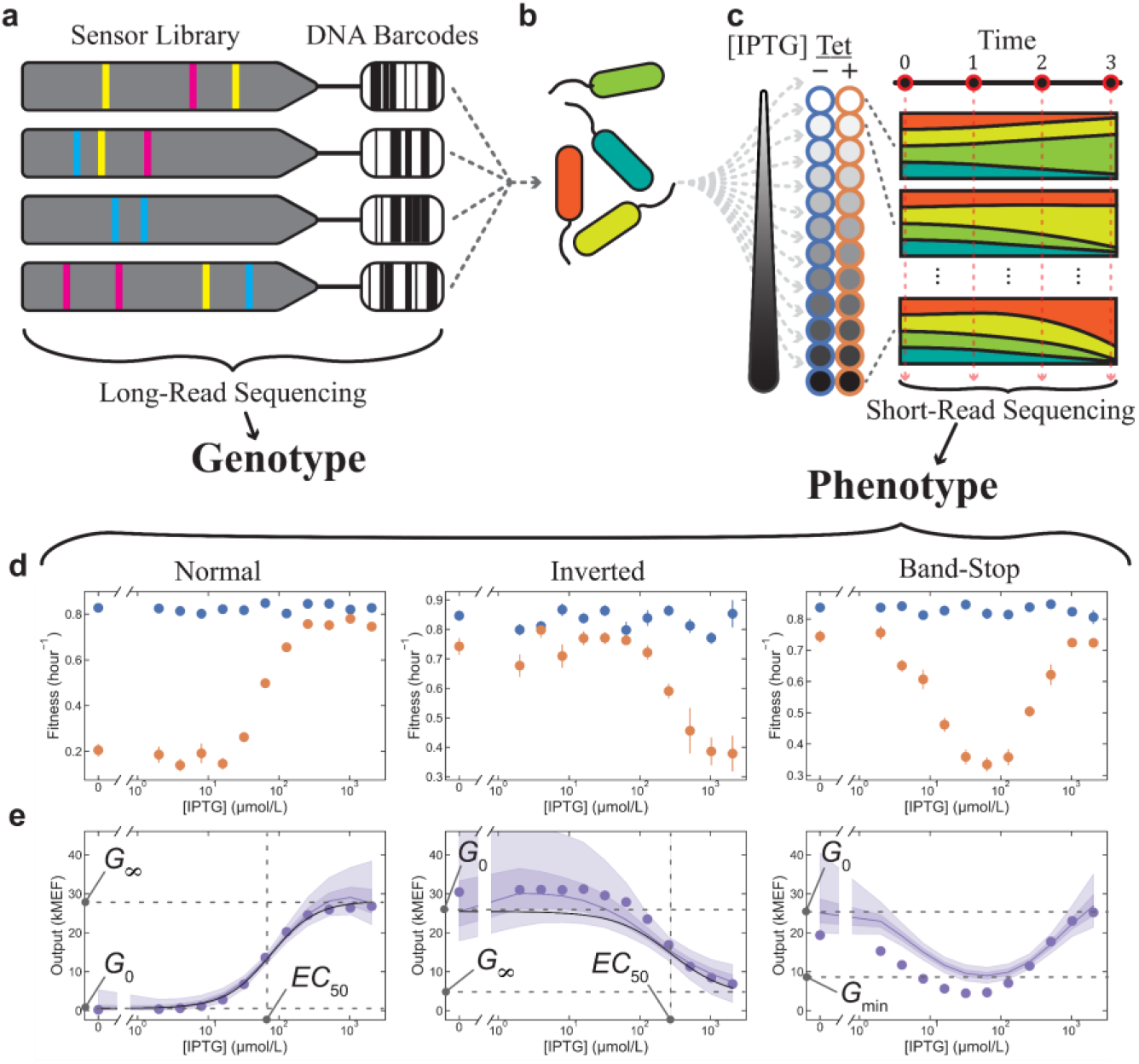
Large-scale allosteric genotype-phenotype landscape measurement. **a**, The sensor library was generated by random mutagenesis of the *lacI* CDS. Each CDS variant was attached to a DNA barcode and inserted into a plasmid where the sensor regulated expression of a tetracycline resistance gene (Extended Data Fig. 8). Long-read sequencing was used to determine the CDS and corresponding DNA barcode for each sensor. **b,** The library was transformed into *E. coli*. **c,** Cells containing the library were grown in 24 chemical environments (12 IPTG concentrations, each with (orange) and without (blue) tetracycline). Cultures were periodically diluted to maintain exponential growth. Short-read sequencing of DNA barcodes at four timepoints was used to measure changes in the relative abundance of each sensor, and those changes were used to determine the fitness associated with each sensor in each environment. **d,** The fitness without tetracycline (blue) is independent of IPTG concentration. The fitness with tetracycline (orange) depends on the IPTG concentration via the dose-response of each sensor. **e,** Dose-response curves for 62 472 sensors were estimated from the fitness measurements with Bayesian inference using a Hill equation model (black lines for normal and inverted sensors) and a Gaussian process (GP) model (purple lines, shaded regions indicate 50 % and 90 % credible intervals). Flow cytometry verification measurements (purple points) generally agreed with Bayesian inference results and also verified the existence of the unexpected band-stop phenotypes. Dose-response output is reported as molecules of equivalent fluorophore (MEF). Error bars indicate ± one standard deviation and are often within markers.

To map the genotype-phenotype landscape for the allosteric LacI sensor, we engineered a genetic construct in which LacI modulates cellular fitness by regulating the expression of a tetracycline resistance gene in response to the concentration of a ligand (isopropyl-β-D-thiogalactopyranoside, IPTG). We then created a library of sensors through mutagenic PCR of *lacI.* To ensure that most sensors in the library could regulate gene expression, we used fluorescence-activated cell sorting (FACS) to enrich the library for sensors with low *G*_0_. Then, using high-accuracy long-read sequencing^15^, we determined the coding DNA sequence (CDS) of every sensor in the library and indexed each CDS to an attached DNA barcode (Fig. 1a).

To characterize the dose-response curve of each sensor in the library, we grew *E. coli* containing the library in 24 chemical environments (12 ligand concentrations, each with and without tetracycline, Fig. 1b-c). We used short-read sequencing of the DNA barcodes to measure the relative abundance of each sensor at four timepoints during growth. We then used the changes in relative abundance to determine the fitness associated with each sensor in each environment. For each sensor in the library, we used the fitness difference (with vs. without tetracycline) from all 12 ligand concentrations to quantitatively determine the dose-response curve using Bayesian inference (Fig. 1d-e, Fig. 2c-f). Most sensors had sigmoidal dose-response curves (e.g. Fig. 2g), which we quantitively characterized by fitting with the Hill equation.

**Fig. 2.**
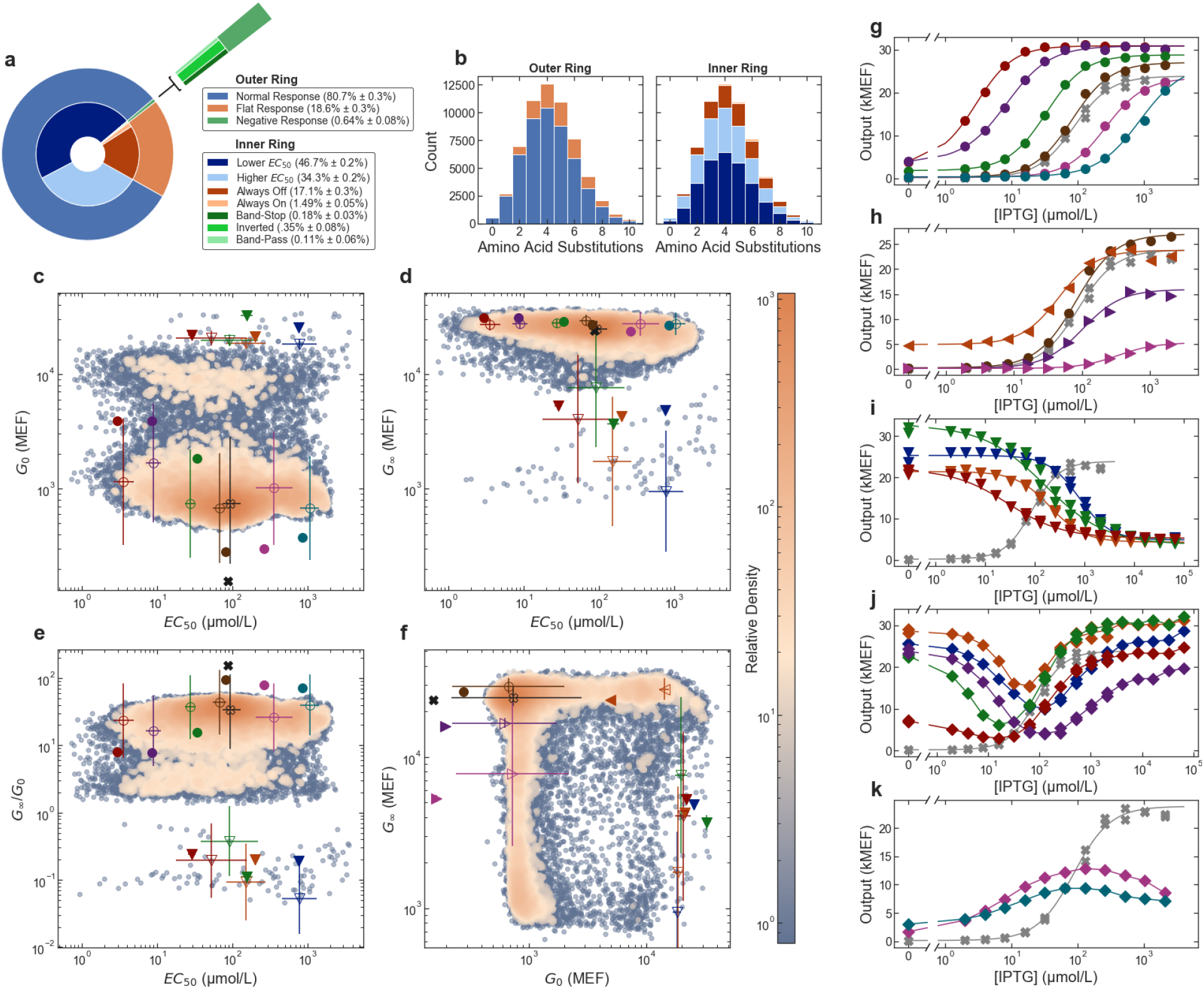
Diversity of sensor phenotypes within the sensor library. **a,b,** Proportion of sensor phenotypes in the library (**a**) and as a function of the number of amino acid substitutions (**b**). The plots in **a** and **b** share the same legend. The outer ring of **a** and the left panel of **b** show the proportion of sensors with dose-response curves that are normal (similar to wild-type), flat-response, or with negative response. The inner ring of **a** and the right panel of **b** show more detailed descriptors of sensors phenotypes. **c-f,**2D plots of Hill equation parameters for all sensors in the library. Each colormap indicates the relative density of sensors within the measured phenotype space. Points plotted with error bars show results for sensors with matching flow cytometry data in **g-i**, with a black ‘X’ indicating the wild type. Open symbols are results from barcode sequencing; solid symbols are results from flow cytometry. In **c-e**, *EC*_50_ values for the barcode sequencing results for normal-response sensors (open circles) are corrected for a slight systematic bias for comparison with flow cytometry results (filled circles, see Extended Data Fig. 5). **g-k,** Dose-response curves measured using flow cytometry to verify sensor phenotypes, including sensors engineered with defined *EC*_50_ (**g**) and *G*_0_ or *G*_∞_ (**h**). Several inverted sensors (**i**) and novel sensor phenotypes, including band-stop (**j**) and band-pass (**k**) sensors were also verified. In all plots, *G*_0_ and *G*_∞_ are given in molecules of equivalent fluorophore (MEF), and the error bars indicate ± one standard deviation.

The sensor library contains 62 472 different CDSs, with an average of 7.0 single nucleotide polymorphisms (SNPs) per CDS. Many SNPs are synonymous, so the library encodes 60 398 different amino acid sequences with an average of 4.4 amino acid substitutions per sensor (Fig. 2b). Synonymous SNPs do not measurably impact the sensor phenotype. So, for subsequent analysis, we considered only amino acid substitutions.

### DNN predicts allosteric dose-response

Our landscape provides the first quantitative dataset of sufficient size to train a DNN capable of predicting dose-response curves for an allosteric genetic sensor. To build the DNN, we adapted a recurrent architecture that captures the context-dependent nature of substitutions on protein function. This architecture predicts Hill equation parameters more accurately than other architectures previously used to predict protein function^9^ (Extended Data Fig. 1). The DNN accurately predicts *EC*_50_ and *G*_∞_(Fig. 3), Hill equation parameters with low measurement uncertainty, but predicts *G*_0_ less accurately, due to larger proportional measurement uncertainty and the small number of measured sensors with substantial changes to *G*_0_ (Fig. 4). For most sensors, the parameter *n* does not vary beyond the measurement uncertainty, so it is not included in the model predictions (Extended Data Fig. 2).

**Fig. 3.**
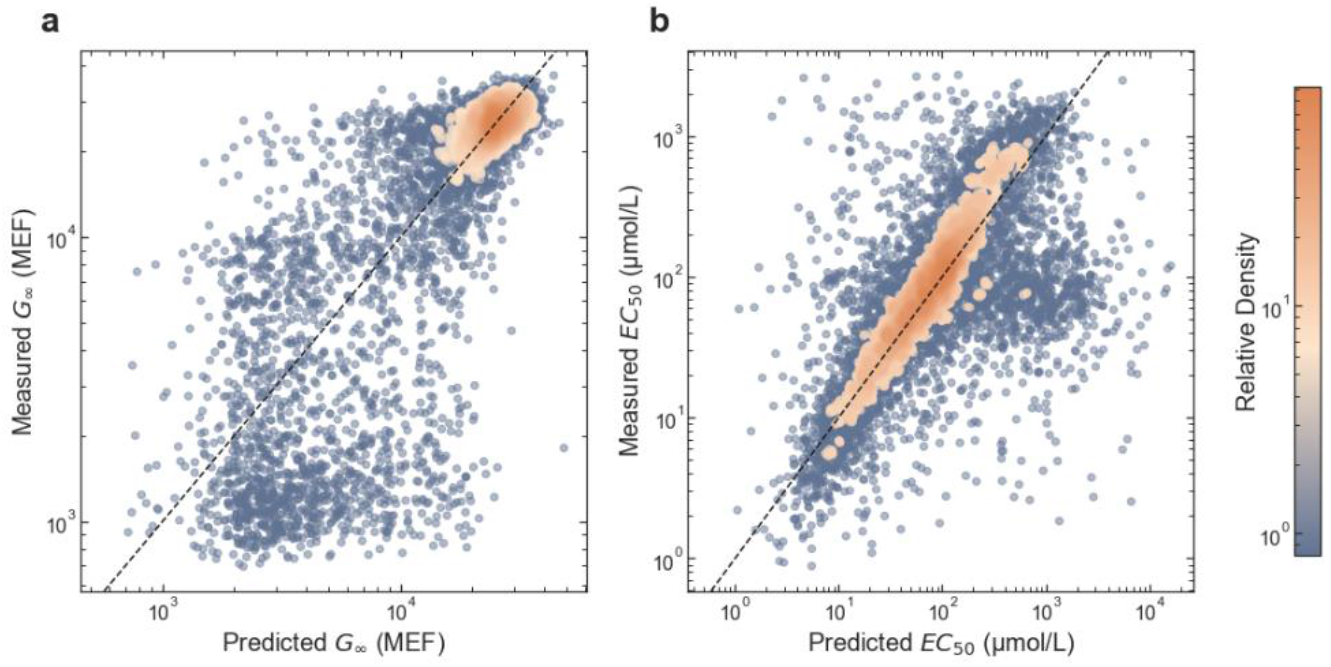
Prediction of sensor phenotypes with recurrent deep neural network (DNN) model. Measured vs. predicted values for Hill equation parameters *G*_∞_ (**a**) and *EC*_50_ (**b**). The plotted results only show holdout data not used in model training. Predictions are taken from the variational posterior mean for each Hill equation parameter. Measured values are the posterior medians from the Bayesian fits to the experimental data. In **a**, *G*_∞_ is given in molecules of equivalent fluorophore (MEF). In **b**, the cluster of points with measured *EC*_50_ near 100 μmol/L and predicted *EC*_50_ near 1000 μmol/L correspond to sensors for which the DNN model provided better *EC*_50_ and *G*_∞_ estimates than the measurement, because *EC*_50_ was near or above the maximum ligand concentration used. For these points, the nearly constant dose-response across the measured concentration range resulted in a large uncertainty for *EC*_50_ and a posterior median near the median of the *EC*_50_ prior (100 μmol/L).

**Fig. 4.**
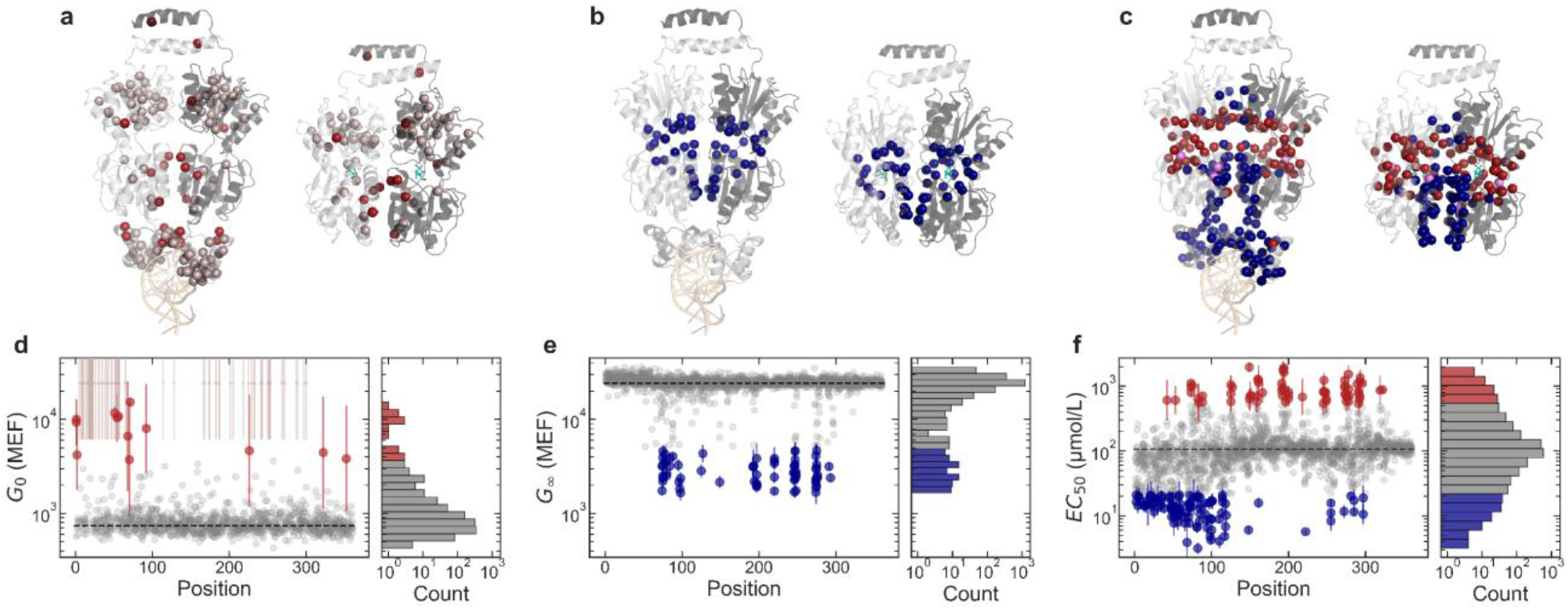
Impact of substitutions on allosteric function. **a-c,** Protein structures showing the locations of substitutions that affect each Hill equation parameter: *G*_0_ (**a**), *G*_∞_ (**b**), *EC*_50_ (**c**). For each, the DNA-binding configuration is shown on the left (DNA in light orange, PDB ID: 1LBG) and the ligand-binding configuration is shown on the right (IPTG in cyan, PDB ID: 1LBH). Both configurations are shown with the view oriented along the protein dimer interface, with one monomer in light gray and the other monomer in dark gray. Colored spheres highlight residues where substitutions cause a greater than 5-fold change in the Hill equation parameter relative to the wild-type. Red spheres indicate residues where substitutions increase the parameter, and blue spheres indicate residues where substitutions decrease the parameter. At three residues (A82, I83, and F161), some substitutions decrease *EC*_50_, while other substitutions increase *EC*_50_ (violet spheres in **c**). **d-f,** Effect of substitutions on *G*_0_ (**d**), *G*_∞_ (**e**), *EC*_50_ (**f**). Scatter plots show the effect of each substitution as a function of position. Substitutions that change the parameter by less than 5-fold are shown as gray points. Substitutions that change the parameter by more than 5-fold are shown as red or blue points with error bars. In **a** and **d**, gray-pink spheres and points indicate positions for substitutions shown by previous work to result in constitutively high *G*(*L*)^18,19^. Histograms to the right of each scatter plot show the overall distribution of substitution effects. *G*_0_ and *G*_∞_ are given in molecules of equivalent fluorophore (MEF). Error bars indicate ± one standard deviation.

Uncertainty quantification is crucial for assessing the confidence of model predictions^16^. Previous examples of DNNs for genotype to phenotype prediction relied on point estimates alone. Here, we quantified uncertainties for model predictions by performing approximate Bayesian inference of the model parameters using a variational method^17^ (Extended Data Fig. 1). The resulting uncertainties allowed us to confidently integrate the experimental and DNN results, and thereby explore a larger landscape than was measured with the experimental data alone.

Specifically, by integrating information about the causal substitutions from multiple genetic backgrounds, the model provides improved estimates of *EC*_50_ and *G*_∞_ for sensors with *EC*_50_ near or above the maximum ligand concentration measured (Fig. 3). In addition, the model confidently predicts *EC*_50_ and *G*_∞_ for single substitutions that are present within the library only in combination with other substitutions (Extended Data Fig. 1). Thereby, we were able to use a sufficiently high mutation rate to explore a broad genotype-phenotype space, while still acquiring the single-substitution information most useful for building quantitative biophysical models of protein function^6–8^.

### Impact of substitutions on allostery

To map the broadly distributed intramolecular interactions that determine the allosteric behavior of LacI, we examined the dose-response curves of sensors with single substitutions. We used experimental data and DNN predictions to determine the dose-response curves for 94 % of the SNP-accessible substitutions (1991 of 2110; 964 directly from measured data, and 1027 from DNN predictions). Most of the 119 substitutions missing from our dataset were probably excluded by FACS during library preparation because they caused a substantial increase in *G*_0_. These include 83 substitutions, located primarily in the DNA-binding domain or at buried residues in the C-terminal core domain, that have been shown to result in constitutively high *G*(*L*)^18,19^. Of the 1991 substitutions included in our dataset, 38 % measurably affect the dose-response curve (beyond a 95 % confidence bound), and the impact of each substitution depends strongly on its location within the protein structure (Fig. 4).

Substitutions that increase *G*_0_ by more than 5-fold but were not excluded by FACS are located either in helix 4 of the DNA-binding domain, along the dimer interface, in the tetramerization helix, or at the protein start codon (Fig. 4a,d). *G*_0_ quantifies the sensor state in the absence of ligand. So, apart from substitutions at the start codon that reduce the number of LacI proteins per cell^20^, these substitutions probably affect either the DNA binding affinity, the allosteric constant of the sensor, or both^8^. Interestingly, substitutions in helix 4 (R51C, Q54K, and L56M) and near the dimer interface (T68N, L71Q) that increase *G*_0_ also decrease *EC*_50_ approximately 10-fold, consistent with a change in the allosteric constant^8^.

Substitutions that decrease *G*_∞_ by more than 5-fold are all located near the ligand-binding pocket or along the dimer interface (Fig. 4b,e). Six of these substitutions also increase *EC*_50_ more than 5-fold (A75T, D88N, S193L, Q248R, D275Y, and F293Y). Except for D88N, which is at the dimer interface, these substitutions are in the ligand-binding pocket. Substitutions near the ligand-binding pocket probably change ligand binding affinity, though previous work has shown that they can also change the allosteric constant ^8^.

In addition to specific substitutions that affect both *G*_∞_ and *EC*_50_, we identified nine positions (N125, P127, D149, V192, A194, A245, N246, T276, Q291), where different substitutions either reduce *G*_∞_ by more than 5-fold or increase *EC*_50_ by more than 5-fold, but not both. All these positions are in the ligand-binding pocket. We also identified five positions (H74, V80, K84, S97, M98) where different substitutions reduce either *G*_∞_ or *EC*_50_ by more than 5-fold but not both (Extended Data Fig. 3). All of these positions are located at the dimer interface.

Substitutions that affect *EC*_50_ are the most numerous and are spread throughout the protein structure, with approximately 9 % and 27 % of all substitutions causing a greater than 5-fold or 2-fold shift in *EC*_50_, respectively (Fig. 4c,f). The strongest effects are from substitutions in the DNA-binding domain, ligand-binding pocket, core-pivot domain, or dimer interface. Substitutions to the DNA-binding domain or dimer interface generally decrease *EC*_50_. Substitutions to the ligand-binding pocket or core-pivot domain generally increase *EC*_50_.

Combining multiple substitutions in a single protein almost always has a log-additive effect on *EC*_50_. Only 0.57 % (12 of 2101) of double substitutions have an *EC*_50_ that differs from the log-additive effects of the single substitutions by more than 2.5-fold (Extended Data Fig. 4). This result, combined with the wide distribution of residues that affect *EC*_50_, suggests that LacI allostery is controlled by a free energy balance with additive contributions from many residues and interactions^7,8,21,22^.

### Precise engineering of genetic sensors

The sensor library contained a wide diversity of sensor function, with *EC*_50_ values spanning more than three orders of magnitude (less than 1 μmol/L to over 1000 μmol/L, Fig. 2c-e) and *G*_0_ and *G*_∞_ values covering more than a 35-fold range (Fig. 2f). To take advantage of this diversity, we demonstrated a new approach to engineering biological function by choosing the corresponding CDS for any desired dose-response curve within the range of the library.

For example, we identified sensors with *EC*_50_ ranging from 3 μmol/L to over 1000 μmol/L (and *G*_0_, *G*_∞_ near the wild-type values). We then verified the dose-response curve of each identified sensor by re-synthesizing the CDS, integrating it into a different genetic construct where it regulated the expression of a fluorescent protein, and measuring fluorescence at 12 ligand concentrations using flow cytometry (Fig. 2g). Comparison between the flow cytometry and the sequencing-based results indicates that we can use this approach to engineer sensors with defined *EC*_50_ within 1.25-fold of a targeted value (Extended Data Fig. 5). In addition, we identified and verified the dose-response curves of sensors with near-wild-type *EC*_50_, but non-wild-type values for *G*_0_ or *G*_∞_ (Fig. 2h). Because of the non-linearity of the fitness impact of tetracycline, the accuracy for engineering sensors with different *G*_0_ and *G*_∞_ levels depends on the growth conditions, particularly the tetracycline concentration. Here, the relative accuracy varied from 1.12-fold near the wild-type *G*_∞_ level to 4.3-fold near the wild-type *G*_0_ level. To our knowledge, this is the first demonstration of engineering allosteric function with quantitatively targeted performance parameters and associated quantification of accuracy.

To further illustrate the range of sensor phenotypes that can be engineered with this approach, we identified and verified the dose-response curves of inverted sensors, i.e. sensors with *G*_0_ > *G*_∞_. (Fig. 2i). Approximately 230 sensors in the library are inverted (0.35 % of the measured library, Fig. 2a), though less than half have a dynamic range with *G*_0_/*G*_∞_ > 2. By examining a set of 43 strongly inverted sensors (with *G*_0_/*G*_∞_ > 2, *G*_0_ > *G*_∞,*wt*_/2, and *EC*_50_ between 3 μmol/L and 1000 μmol/L), we identified 10 substitutions associated with the inverted phenotype (S70I, K84N, D88Y, V96E, A135T, V192A, G200S, Q248H, Y273H, A343G). However, none of these substitutions are present in more than 12 % of the strongly inverted sensors, and 51 % of the strongly inverted sensors have none of these substitutions. Furthermore, the strongly inverted sensors are more genetically distant from each other than randomly selected sensors from the library (Extended Data Fig. 6).

Inverted sensors can provide specific insight into the allosteric mechanisms of LacI, since they require inversion of both the allosteric constant^6^ and the relative ligand-binding affinity between the two conformations^7^. Although the set of strongly inverted sensors are genetically diverse, many of them share common features that may account for these allosteric changes. First, 67 % of the strongly inverted sensors have substitutions near the ligand-binding pocket (within 7 Å), which likely contribute to the change in ligand-binding affinity. Surprisingly, 21 % of the strongly inverted sensors have no substitutions within 10 Å of the binding pocket, so binding affinity must be indirectly affected by distal substitutions in these sensors. Second, nearly all strongly inverted sensors have substitutions at the dimer interface (91 %, compared to 54 % for the full library), with most (70 %) having substitutions in helix 5 (47 %), helix 11 (28 %), or both (5 %, Extended Data Fig. 6). This suggests that residues in those structural domains play an important role in determining the allosteric constant.

### Novel sensor phenotypes

In addition to the normal and inverted phenotypes, we were surprised to find sensors with dose-response curves that do not match the sigmoidal form of the Hill equation. Specifically, we found sensors with band-pass or band-stop dose-response curves, i.e. sensors that repress or activate gene expression only over a narrow range of ligand concentrations. We verified the dose-response curves of 13 band-stop sensors and three band-pass sensors using flow cytometry (Fig. 2j,k). To our knowledge, this is the first identification of single-protein genetic sensors with band-stop dose-response curves.

Phenotypic similarities between the band-stop and inverted sensors (i.e. high *G*_0_ and initially decreasing gene expression as ligand concentration increases) imply similar biophysical requirements. However, the two-step switching of band-stop sensors suggests the relevant free energy changes may be more entropic than structural^23^. Indeed, substitutions associated with band-stop sensors are remarkably different from those found for the inverted sensors. While inverted sensors often have substitutions near the ligand-binding pocket and dimer interface, a set of 31 strong band-stop sensors are twice as likely as the full library to have substitutions in helix 9 (32 % compared to 16 %) and nearly four times as likely to have substitutions in strand J (13 % compared to 3.4 %). Helix 9 is on the periphery of the protein, and strand J is in the center of the C-terminal core domain. Furthermore, 100 % of the strong band-stop sensors had substitutions in the C-terminal core of the protein, compared with 79 % of the full library (Extended Data Fig. 6).

To further investigate the band-stop phenotype, we identified a band-stop sensor from the landscape measurement with only three substitutions (R195H/G265D/A337D). We synthesized sensors with all possible combinations of those substitutions and measured their dose-response curves. Although each single substitution resulted in a sigmoidal dose-response similar to wild-type LacI, the combination of two substitutions (R195H/G265D) gave rise to the band-stop phenotype (Extended Data Fig. 7). The existence of a band-stop sensor only two substitutions from the wild-type LacI indicates that novel and potentially useful phenotypes exist with readily accessible genotypes. This conclusion is further supported by the relative abundance of band-stop sensors, which comprise nearly 0.2 % of the full library. For comparison, directed evolution to engineer new phenotypes often involves the creation of libraries with more than 10^8^ variants^24^.

## Discussion

Overall, our findings suggest that a surprising diversity of useful and potentially novel allosteric protein phenotypes exist with genotypes that are discoverable via large-scale landscape measurements. As DNA sequencing improves, large-scale landscape measurements like this one will become increasingly accessible and will improve our fundamental understanding of biology while also enabling robust and scalable engineering of biological function for applications such as living therapeutics^25^, structured materials fabrication^26^, and engineered morphogenesis^27^.

## Accession codes

### Primary accession codes

#### Bioproject

##### Long-Read and Barcode Sequencing

PRJNA643436

#### Genbank

##### Library Plasmid

MT702633

##### Verification Plasmid

MT702634

### Referenced accession codes

#### Protein Data Bank

##### DNA-binding conformation of LacI

1LBG

##### Ligand-binding conformation of LacI

1LBH

## Extended Data Figures and captions

**Extended Data Fig. 1.**
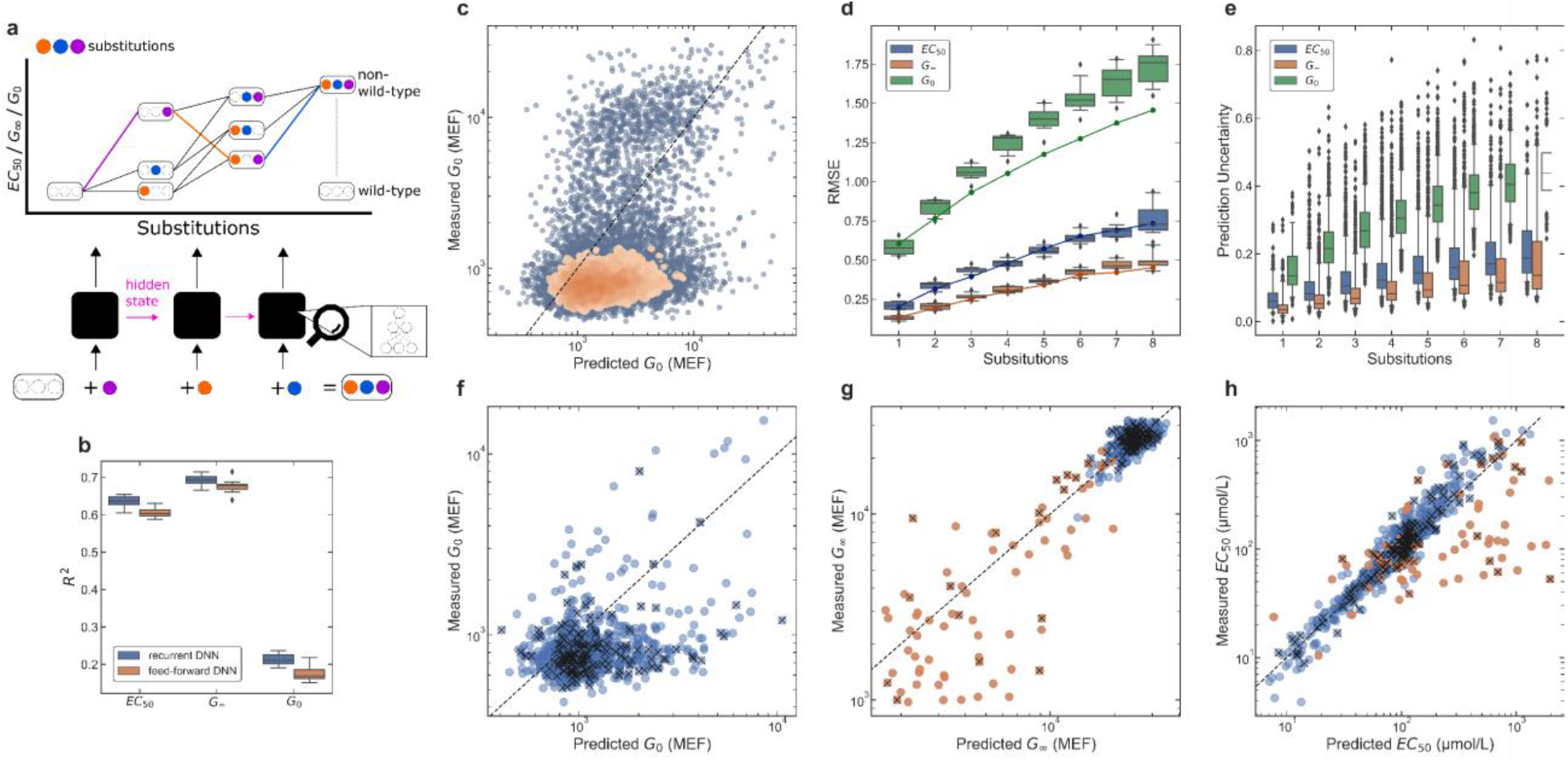
Deep neural network (DNN) model evaluation. **a**, Schematic illustration of recurrent deep neural network model architecture. Using the wild-type sequence as a starting point, the model predicts the Hill equation parameters for non-wild-type sequences by composing the individual parameter changes due to sequential substitutions along a mutational path. These changes are then added together to predict the final parameter values. All paths to a non-wild-type sequence converge to the same value, and this fact is leveraged to build a recurrent neural network model that learns to predict the individual substitution effects, given previous substitutions. Potential non-additive effects are captured by the hidden state of the model, which predicts the change in parameter value for the most recent substitution and serves as a set of latent variables for predicting subsequent substitutions. Note that sensors with intermediate sequences may be present in the library, but this is not necessary to train the model. The model will still learn to predict intermediate steps in the path, even if that data is not present. **b**, Performance of recurrent and feed-forward DNN models. The boxplot summarizes the *R*^2^ values for ten cross-validation tests for each model. The recurrent DNN model generally outperforms the feed-forward model, giving higher *R*^2^ values for each of the Hill equation parameters. **c**, Measured vs. predicted values for Hill equation parameter *G*_0_ are plotted with a colormap indicating the relative density of data. Predictions were taken from the variational posterior mean for *G*_0_. Results are plotted only for the holdout data not used in model training. **d**, Model prediction error for each Hill equation parameter as a function of the number of substitutions from the wild-type sequence. Prediction error is shown as the root-mean-square error (RMSE) for the base-ten logarithm of each Hill equation parameter. The boxplot shows the distribution of RMSE values from the ten cross-validation tests for the recurrent DNN model. Solid lines show the RMSE for the holdout data not used for training. **e**, Model prediction uncertainty for the base-ten logarithm of each Hill equation parameter as a function of the number of substitutions from the wild-type sequence. The boxplot shows the distribution of posterior standard deviation values for the set of sensors with the indicated number of substitutions. Both the RMSE (**d**) and the prediction uncertainty (**e**) increase with increasing mutational distance from the wild-type sequence. **f-h**, Measured vs. predicted values for the effect of single substitutions on Hill equation parameters *G*_0_ (**f**), *G*_∞_ (**g**), and *EC*_50_ (**h**). Blue symbols show data for all of the single-substitution sensors in the library. Orange symbols show data for sensors with a high uncertainty for the measured *EC*_50_ (std(log_10_(*EC*_50_)) > 0.35). The *EC*_50_ uncertainty for those sensors was relatively high either because *G*_∞_ was similar to *G*_0_ and/or because *EC*_50_ was near or above the maximum ligand concentration used (2048 μmol/L). For those sensors, in the analysis of single-substitution effects, the DNN model result for *G*_∞_ and *EC*_50_ was used in place of the experimental result. Points marked with an ‘x’ were in the holdout data not used for model training. In all plots, *G*_0_ and *G*_∞_ are given in molecules of equivalent fluorophore (MEF).

**Extended Data Fig. 2.**
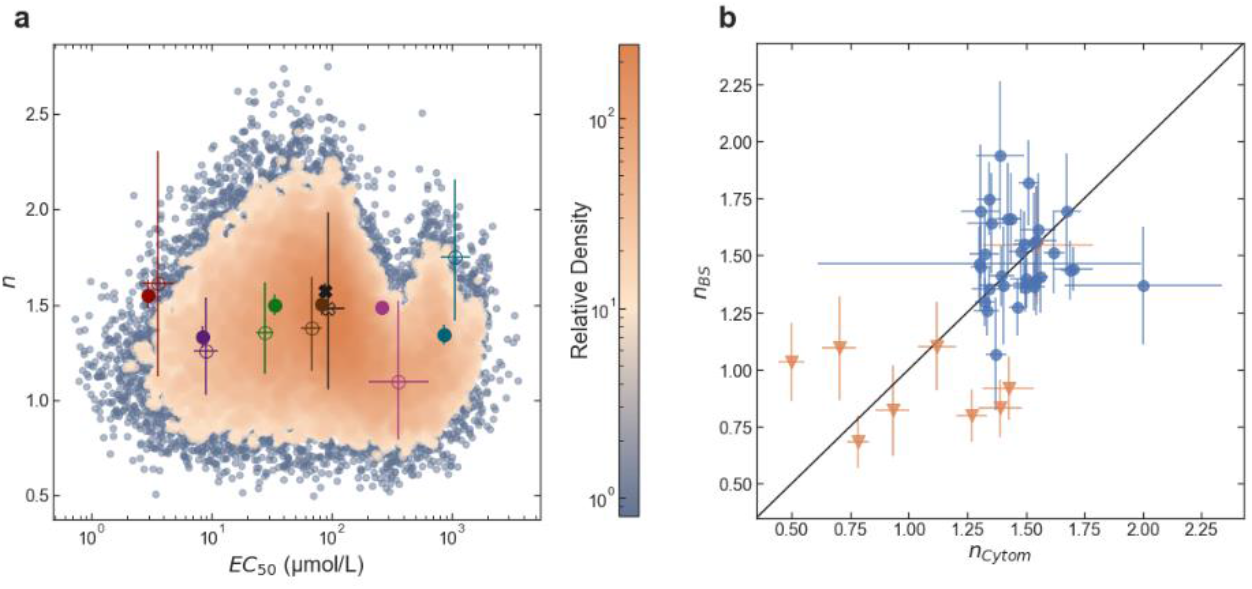
Data for the *n* parameter of the Hill equation. **a,**2D plot of Hill equation *n* parameter vs. *EC*_50_ for all sensors in the library. The density colormap indicates the relative density of sensors within the measured phenotype space. Plotted points show results for the verified phenotypes plotted in Fig. 2g. The wild-type phenotype is marked with a black ‘X’. Open symbols are the results from barcode sequencing; solid symbols are the results from flow cytometry. *EC*_50_ values for the barcode sequencing results (open circles) are corrected for a slight systematic bias for comparison with the flow cytometry results (filled circles, see Extended Data Fig. 5). **b**, Comparison between the *n* parameter from barcode sequencing (*n_BS_*) and from flow cytometry (*n_Cytom_*). Blue circular symbols show data for normal-phenotype sensors (sensors with *G*_0_ and *G*_∞_ near wild-type values) and orange triangular symbols show data for inverted sensors. Inverted sensors generally have a lower *n* parameter than normal phenotype sensors, a trend that is captured by both measurement methods. In both **a** and **b**, the plotted points and error bars indicate the posterior median and standard deviation from the Bayesian inference.

**Extended Data Fig. 3.**
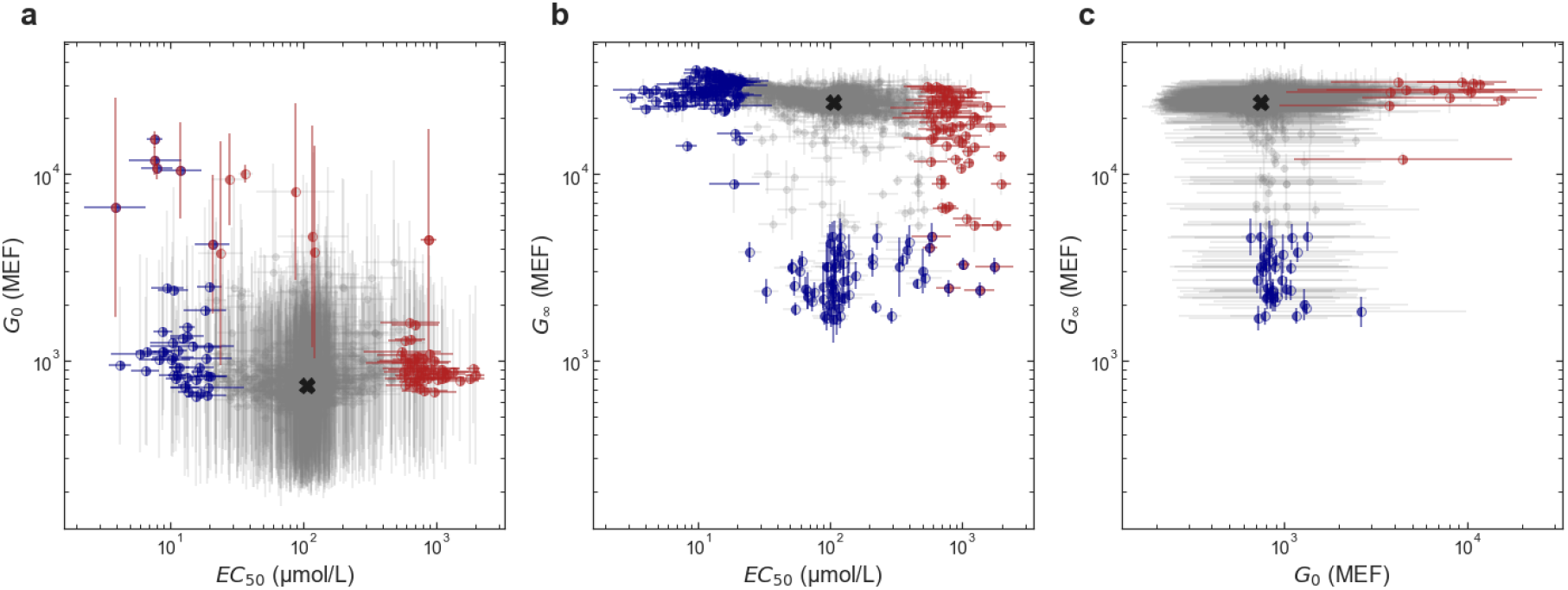
Multiparametric impact of substitutions on allosteric function. The effect of single amino acid substitutions on the Hill equation parameters are plotted to show the joint effect of each substitution on two Hill equation parameters. **a**, *G*_0_ vs. *EC*_50_. **b**, *G*_∞_ vs. *EC*_50_. **c**, *G*_∞_ vs. *G*_0_. In each plot, substitutions that change both Hill equation parameters by less than 5-fold are shown as light gray points, and substitutions that change one or both Hill equation parameters by more than 5-fold are shown as red or blue points with error bars. As in Fig. 4, red indicates a decrease in Hill equation parameter and blue indicates an increase. The left half of each symbol and the y-error bar are colored based on the y-axis parameter; the right half of each symbol and the x-error bars are colored based on the x-axis parameter. The wild-type phenotype is indicated with a black ‘X’ in each plot. In all plots, *G*_0_ and *G*_∞_ are given in molecules of equivalent fluorophore (MEF). Error bars indicate ± one standard deviation.

**Extended Data Fig. 4.**
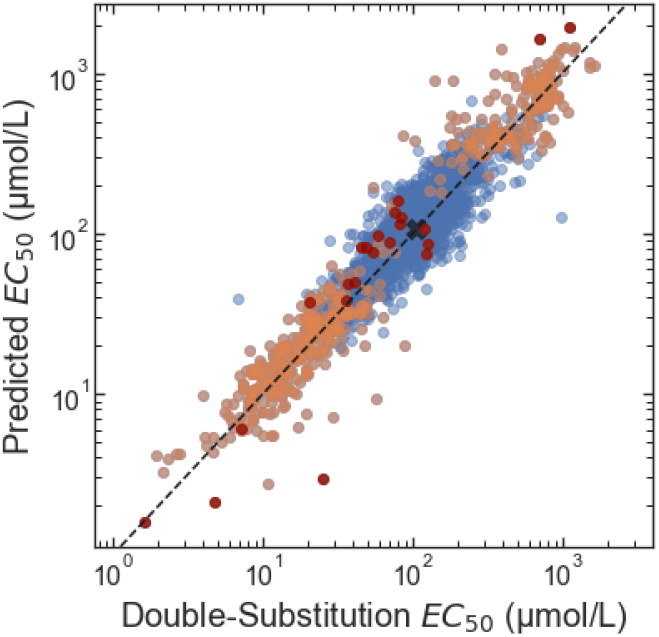
Log-additivity of *EC*_50._ The predicted *EC*_50_ for double substitutions (y-axis) was calculated from single-substitution *EC*_50_ values assuming log-additivity relative to the wild-type *EC*_50_: (*EC*_50,*AB*_ – *EC*_50,*wt*_) = (*EC*_50,*A*_ – *EC*_50,*wt*_) + (*EC*_50,*B*_ – *EC*_50,*wt*_), where ‘*wt*’ indicates the wild-type, ‘*A*’ and ‘*B*’ indicate the single-substitutions, and ‘*AB*’ indicates the double substitution. The actual double-substitution *EC*_50_ (x-axis) is from the experimental measurement (barcode sequencing). Orange points mark double substitutions in which one of the single substitutions causes a greater than 3-fold change in *EC*_50_. Dark red points mark double substitutions in which both single substitutions cause a greater than 3-fold change in *EC*_50_. The wild-type *EC*_50_ is marked with a black ‘X’. For this analysis, only experimental data was used (no results from the DNN model). Also, only data from sensors with low *EC*_50_ uncertainty (std(log_10_(*EC*_50_)) < 0.35) were used.

**Extended Data Fig. 5.**
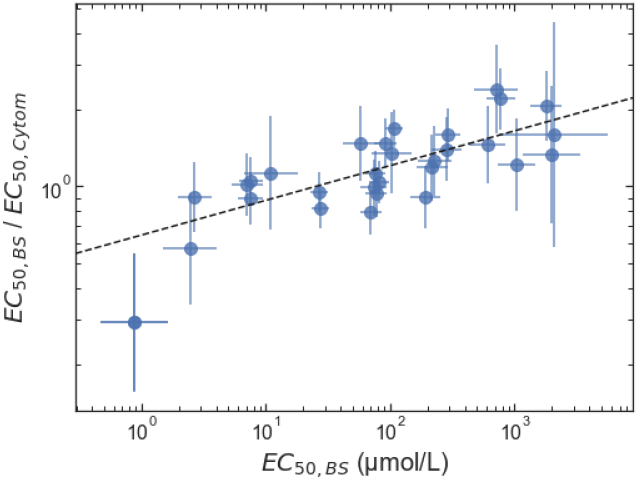
Accuracy of *EC*_50_ from barcode sequencing measurements. The ratio of the *EC*_50_ determined by barcode sequencing (*EC*_50*,BS*_) to the *EC*_50_ determined by cytometry (*EC*_50*,Cytom*_) is plotted vs. *EC*_50*,BS*_ for 31 sensors with *G*_0_ and *G*_∞_ near the wild-type values. For sensors with *EC*_50_ greater than 1 μmol/L, the root-mean-square (RMS) difference between log_10_(*EC*_50*,BS*_) and log_10_(*EC*_50*,Cytom*_) is 0.16, corresponding to an relative accuracy of 1.45-fold (i.e. 10^0.16^). The data show a slight systematic bias, with the barcode sequencing method underestimating *EC*_50_ when it is less than 25 μmol/L and overestimating *EC*_50_ when it is greater than 25 μmol/L. The most accurate results are obtained by correcting the barcode sequencing result based on a fit to this bias (dashed line in the figure). After applying this correction, the RMS difference between log_10_(*EC*_50*,BS*_) and log_10_(*EC*_50*,Cytom*_) is 0.097, corresponding to an relative accuracy of 1.25-fold.

**Extended Data Fig. 6.**
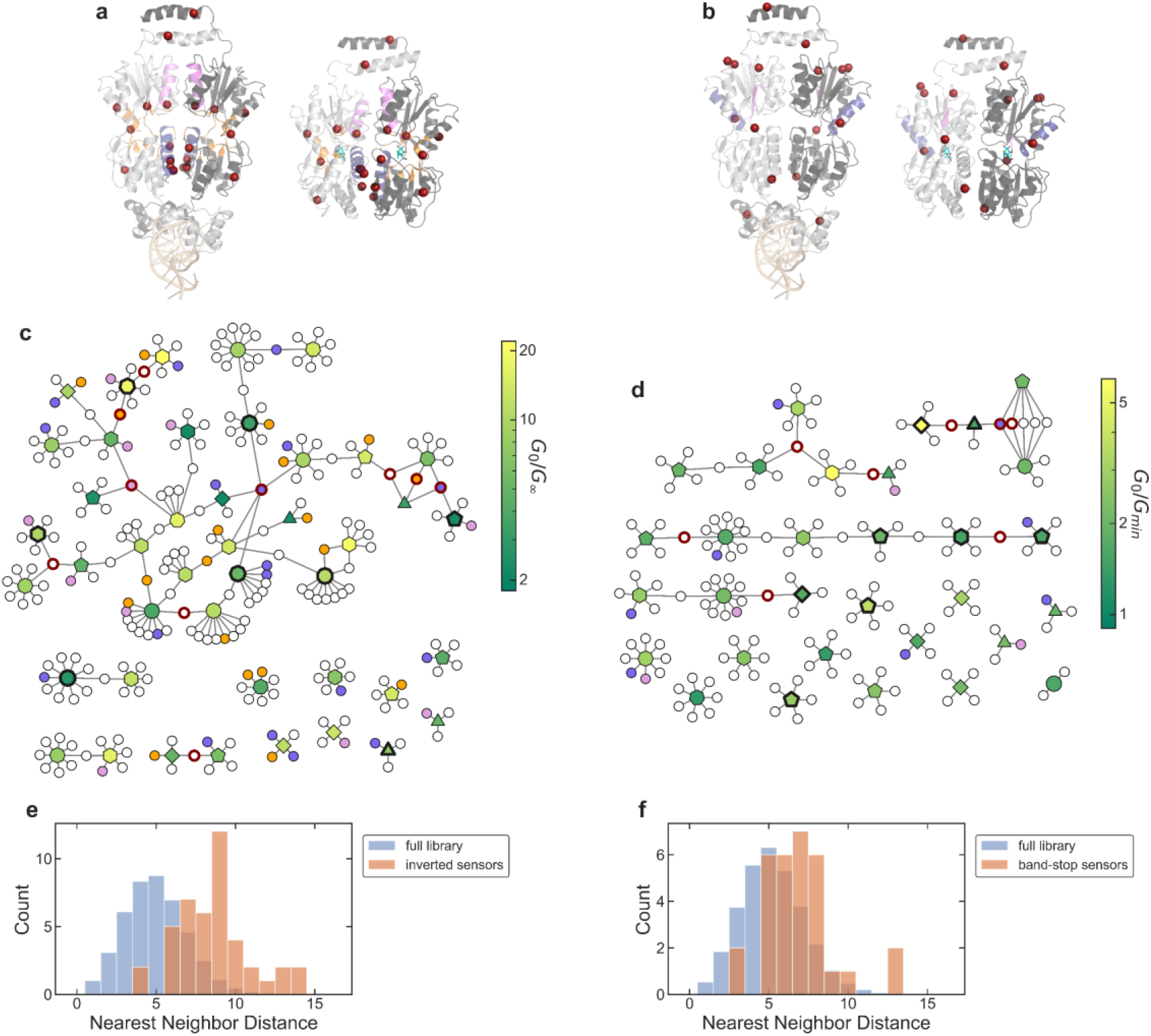
Analysis of inverted and band-stop genotypes. **a,b**, Location of substitutions associated with strongly inverted (**a**) and strong band-stop (**b**) sensors. For each plot, the DNA-binding configuration of LacI is shown on the left (PDB ID: 1LGB), with the DNA operator at the bottom in light orange; the ligand-binding configuration is shown on the right (PDB ID: 1LBH), with IPTG in cyan. Both configurations are shown with the view oriented along the protein dimer interface, with one monomer in light gray and the other monomer in dark gray. The locations of associated (i.e. high-frequency) substitutions are highlighted as red spheres, and structural domains where inverted or band-stop sensors have substitutions at a significantly higher frequency than the full library are shaded with different colors. For strongly inverted sensors (**a**), helix 5 is shaded blue, helix 11 is shaded violet, and the residues near the ligand-binding pocket are shaded orange. For strong band-stop sensors (**b**), helix 9 is shaded blue, and strand J is shaded violet. **c,d**, Network diagrams showing relatedness among genotypes for strongly inverted (**c**) and strong band-stop (**d**) sensors. Within each network diagram, sensors are represented by polygon-shaped nodes, with a colormap indicating the *G*_0_/*G*_∞_ or *G*_0_/*G_min_* ratio (see Fig. 1e). The number of sides of the polygon indicates the number of substitutions relative to the wild-type, and bold outlines indicate sensors that were verified with flow cytometry. Smaller circular nodes represent substitutions, with lines showing the substitutions for each sensor. Bold red outlines on the substitution nodes indicate the associated substitutions shown as spheres in **a-b**, and the shading of substitution nodes matches the shading used to highlight structural domains in **a-b**. **e,f**, Nearest neighbor distance histograms. In each plot, the orange bars show the distribution of nearest neighbor Hamming distance for the amino acid sequences for strongly inverted (**e**) and strong band-stop (**f**) sensors, and the blue bars show the distribution of nearest neighbor Hamming distance for a similar number of randomly selected sequences from the full library. The full-library histograms (blue bars) are averaged over 1000 iterations of randomly selected sequences.

**Extended Data Fig. 7.**
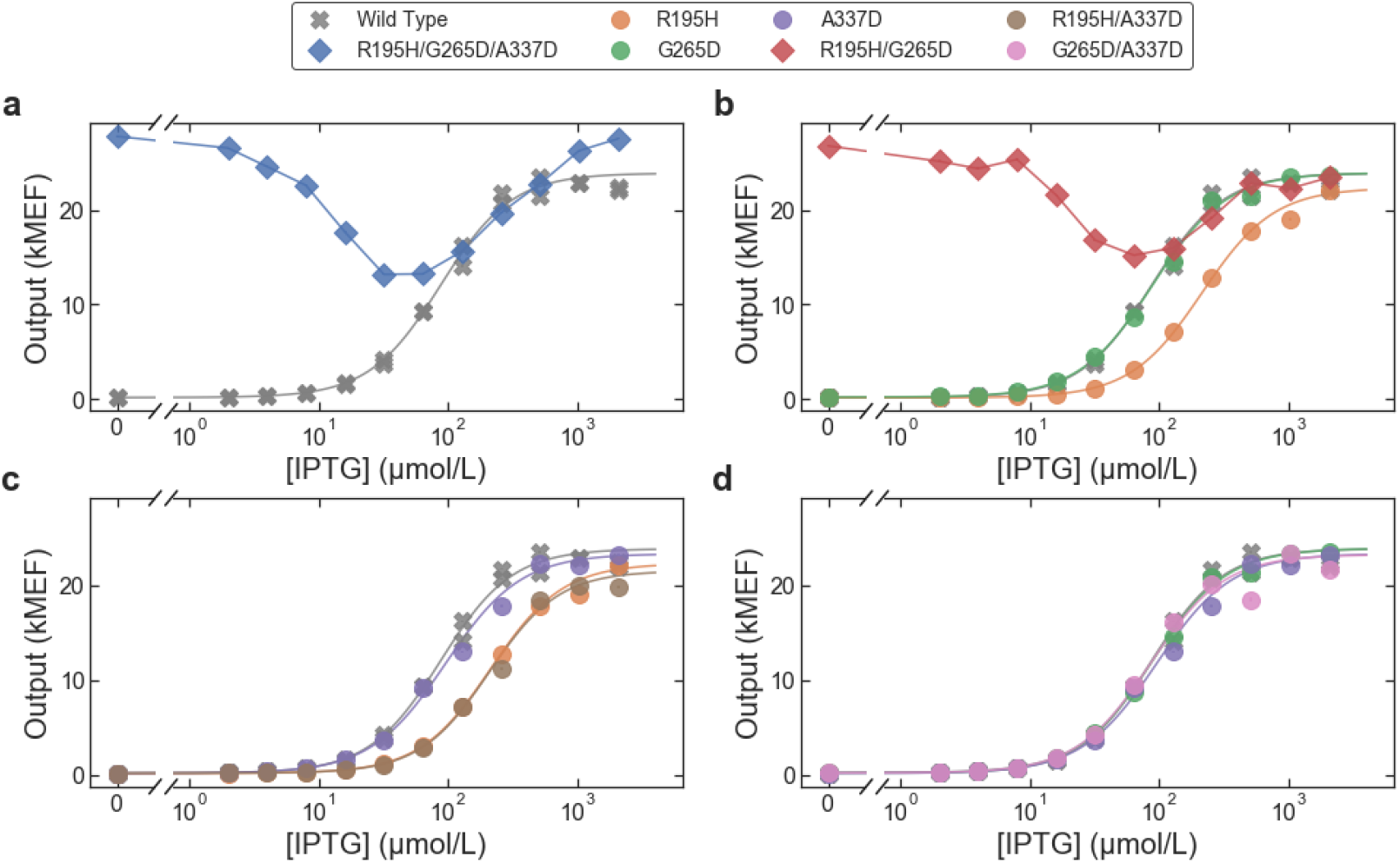
The band-stop phenotype emerges from the combination of two substitutions. **a**, Dose-response curves measured with flow cytometry for the wild-type LacI sensor and a strong band-stop sensor identified in the library with only three substitutions (R195H, G265D, A337D). **b-d,** Dose-response curves for sensors containing single- and double-substitution permutations of the three substitutions in the band-stop sensor. Each plot shows two single substitutions and the combined double substitution. The single substitutions R195H (orange) or G265D (green) result in sigmoidal dose-response curves similar to the wild-type, but the combination of the two, R195H/G265D (red), results in a band-stop phenotype (**b**). All other combinations of single- and double-substitutions result in sigmoidal dose-response curves similar to the wild-type (**c-d**).

**Extended Data Fig. 8.**
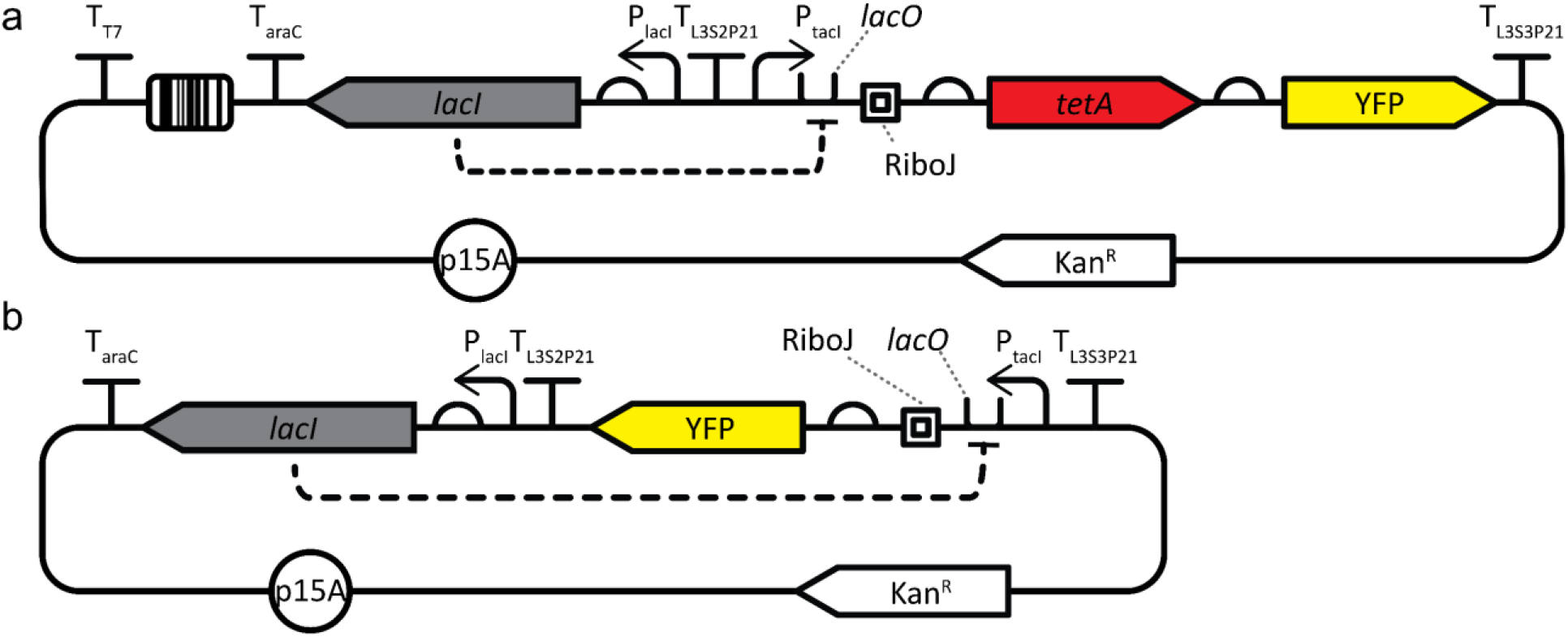
Plasmid maps for the two plasmids used in this work. Both plasmids contained the p15A origin of replication (p15A), kanamycin resistance gene (Kan^R^), and *lacI* CDS. In both plasmids, the encoded LacI protein transcriptionally regulated an output that was driven by the P_tacI_ promoter, *lacO* operator, and RiboJ transcriptional insulator. **a,** The Library Plasmid was used for the barcode sequencing fitness measurements of the sensor library. The sensor library, including sensors and corresponding barcodes, was cloned into the Library Plasmid. Sensors in the Library Plasmid regulated the expression of a tetracycline resistance gene, *tetA*. The Library Plasmid also encoded YFP, which was used during cloning. The expression of *tetA* (and YFP) from the Library Plasmid was transcriptionally terminated with T_L3S3P21_. **b,** The Verification Plasmid was used to verify the dose-response curves of approximately 120 sensors from the library. To verify the dose-response curve of a sensor, the CDS for that sensor was chemically synthesized and cloned into the Verification Plasmid, where the sensor regulated the expression of YFP. The dose-response curve was then measured using flow cytometry.

**Extended Data Fig. 9.**
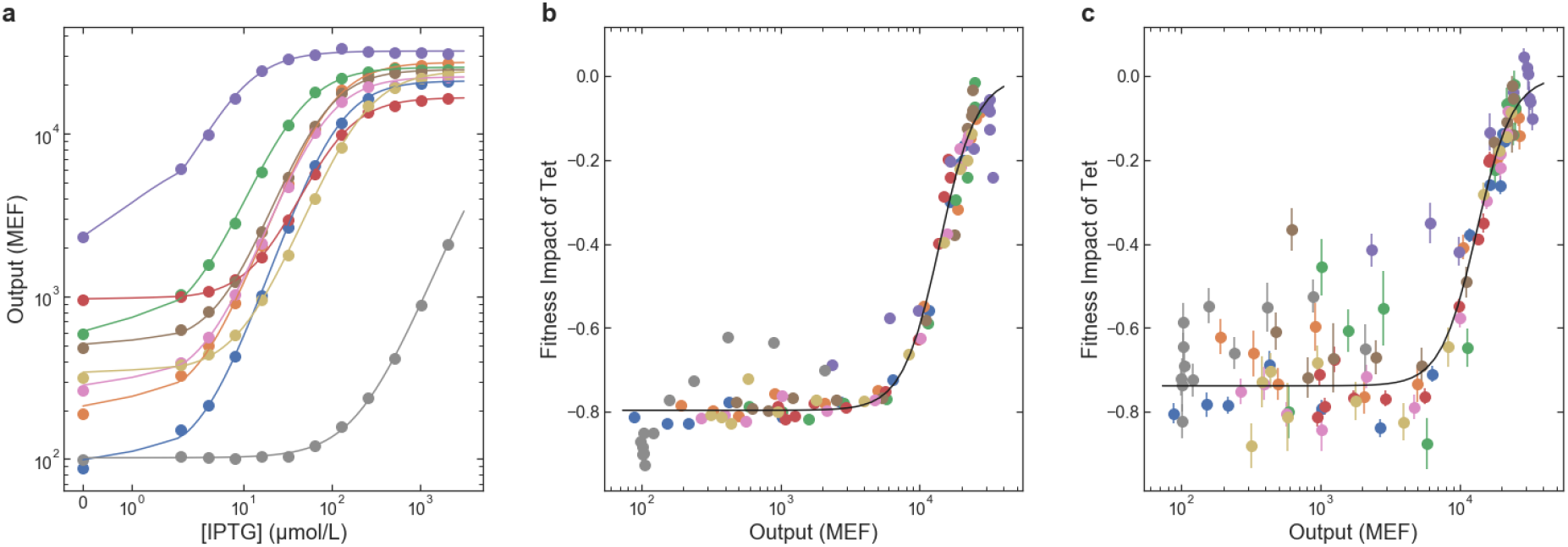
Calibration data for the determination of dose-response curves using barcode sequencing. **a**, Dose-response curves measured with flow cytometry for nine sensors used to calibrate sensor output with fitness. **b**,**c**, The fitness impact of tetracycline (from the barcode sequencing) plotted vs. sensor output (from flow cytometry). The fitness impact of tetracycline is defined as the decrease in fitness (*E. coli* growth rate) measured with tetracycline vs. without tetracycline normalized by the fitness measured without tetracycline, i.e. (*μ^tet^*/*μ*^0^ – 1) from equation (2). **b**, Data from a small-scale test library containing only the calibration sensors. **c,** Data from measurement of the full library. Results for different calibration sensors are shown with different colored symbols. The solid black lines in **b-c** show the results of a fit using equation (2). Error bars indicate ± one standard deviation. The results for the full library measurement are noisier than the small-scale test because of the lower read count per sensor (approximately 100-fold difference), but the general trend and the fits for both results are similar.

## Methods

### Strain, plasmid, and library construction

All reported measurements were completed using *E. coli* strain MG1655Δ*lac* (described previously)^28^. Briefly, strain MG1655Δ*lac* was constructed by replacing the lactose operon of *E. coli* strain MG1655 (ATCC #47076) with the bleomycin resistance gene from *Streptoalloteichus hindustanus* (*Shble*).

Two plasmids were used for this work: a Library Plasmid (Extended Data Fig. 8a) used for the measurement of the genotype and phenotype of the entire library, and a Verification Plasmid (Extended Data Fig. 8b) used to verify the function of 120 sensors from the library chosen to test the accuracy of the landscape measurement and to demonstrate the range of sensor function.

The Library Plasmid contained the *lacI* CDS and the lactose operator (*lacO*) regulating the transcription of a tetracycline resistance gene, *tetA*, which, in the presence of tetracycline, confers a measurable change in fitness connected with the sensor output. The Library Plasmid also encoded Enhanced Yellow Fluorescent Protein (YFP), which was used to select a starting library of sensors with low gene expression in the absence of IPTG using FACS (Sony SH800S Cell Sorter).

The Verification Plasmid contained a similar system in which *lacI* and *lacO* regulate the transcription of only YFP. The Verification Plasmid was used to measure dose-response curve of each sensor using flow cytometry. Each sensor chosen from the library for verification was chemically synthesized (Twist Biosciences), inserted into the Verification Plasmid, and transformed into *E. coli* strain MG1655Δ*lac* for flow cytometry measurements to confirm the dose-response curve inferred from the barcode sequencing measurements.

The sensor library was generated by error-prone PCR of the wild-type *lacI* CDS from the genome of MG1655. The library was inserted into the Library Plasmid along with randomly synthesized DNA barcodes. Each barcode consisted of 54 random nucleotides introduced with PCR primers (Integrated DNA Technologies). Most of the sensors in the initial library had high *G*(0), i.e. the *I*^−^ phenotype^18^. To generate a library of mostly functional allosteric sensors, we used fluorescence activated cell sorting (FACS) to select a portion of the library with low fluorescence in the absence of ligand (Sony SH800S Cell Sorter). To allow comprehensive long-read sequencing of the library (PacBio sequel II, see below), we further reduced the library size by dilution of the FACS-selected library to create a population bottleneck of the desired size. For the work reported here, we used a library of approximately 2 × 10^5^ sensors (determined by serial plating and colony counting).

### *E. coli* culture conditions

*E. coli* cultures were grown in a rich M9 media comprising M9 salts (3 g/L KH_2_PO_4_, 6.78 g/L Na_2_HPO_4_, 0.5 g/L NaCl, 1 g/L NH_4_Cl) supplemented with 0.1 mmol/L CaCl_2_, 2 mmol/L MgSO_4_, 4 % glycerol, and 20 g/L casamino acids. When required for plasmid maintenance, media was supplemented with 50 μg/mL kanamycin.

A fully automated microbial growth and measurement system facilitated the simultaneous measurement of the library across 24 chemical environments. This system prepared all growth conditions, including the addition of IPTG (ligand) and tetracycline. Bacterial cultures were grown in 0.5 mL of media in clear-bottom polystyrene 96-well plates with 1.1 mL volume square wells (4titude, 4ti-0255). Growth plates were sealed with a gas permeable membrane (4titude, 4ti-0598) and incubated at 37 °C (with a 1 °C gradient to minimize condensation) while shaking at 807 double-orbital cycles per minute in a plate reader (BioTek, Neo2SM). Optical density at 600 nm (OD_600_) was measured every 5 minutes during growth.

### Growth protocol for landscape measurement

To begin the LacI genotype-phenotype landscape measurement, a culture of *E. coli* containing the sensor library was mixed at a 99:1 ratio with a culture of an *E. coli* spike-in control with known fitness (see below). The culture was loaded into the automated microbial growth and measurement system where it was distributed across a 96-well plate and then grown to stationary phase (12 hours). Cultures were then diluted 50-fold into a new 96-well plate, Growth Plate 1, containing 11 rows with a 2-fold serial dilution gradient of IPTG with concentrations ranging from 2 μmol/L to 2048 μmol/L plus one row without IPTG. Growth in IPTG allowed the sensors to reach a steady state, including basal expression of tetA in each IPTG concentration. Growth Plate 1 was grown for 160 minutes, corresponding to approximately 3.3 generations, and then diluted 10-fold into Growth Plate 2. Growth Plate 2 contained the same IPTG gradient as Growth Plate 1 with the addition of tetracycline (20 μg/mL) to half of the wells, resulting in 24 chemical environments, with 4 duplicate wells for each environment. Growth Plate 2 was grown for 160 minutes and then diluted 10-fold into Growth Plate 3, which contained the same 24 chemical environments as Growth Plate 2. This process was repeated for Growth Plate 4, which also contained the same 24 chemical environments. The total growth time for the fitness measurements in the 24 chemical environments, 480 minutes across Growth Plates 2-4, corresponded to approximately 10 generations for the fastest-growing cultures. The 50-fold dilution factor from stationary phase into Growth Plate 1 and the 160 minute growth time per plate were chosen to maintain the cultures in exponential growth for the entire 480 minutes.

After each growth plate was used to seed the subsequent plate (or at the end of 160 minutes for Growth Plate 4), the remaining culture volumes for each chemical environment (approximately 450 μL/well, four duplicates per plate) were combined and pelleted by centrifugation (3878 *g* for 10 minutes at 23 °C). Plasmid DNA was then extracted with a custom method using reagents from the QIAprep Miniprep Kit (Qiagen cat. #27104) on an automated liquid handler equipped with a positive-pressure filter press.

### Barcode sequencing

After plasmid extraction, each set of 24 plasmid DNA samples was prepared for barcode sequencing using a custom sequencing sample preparation method on a second automated liquid handler. Briefly, the plasmid DNA was linearized with ApaI restriction enzyme. Then, a 3-cycle PCR was performed to attach sample multiplexing tags to the resulting amplicons so the different samples could be distinguished when pooled and run on the same sequencing flow cell. Eight forward index primers and 12 reverse index primers were used to label the amplicons from each sample across the 24 chemical environments and the four time points. After a magnetic-bead-based cleanup step, a second, 15-cylce PCR was run to attach the standard Illumina paired-end adapter sequences and to amplify the resulting amplicons for sequencing. After a second magnetic-bead-based cleanup, the 24 samples from each time point were pooled and stored at 4 °C until sequencing. For sequencing, DNA was diluted to a final concentration of approximately 5 nmol/L and combined with 20 % phiX control DNA. DNA from each of the 4 time points was sequenced in a separate lane on an Illumina HiSeqX using paired-end mode with 150 bp in each direction.

To count DNA barcodes and estimate the fitness associated with each sensor, the sequencing data was analyzed using custom software written in C# and Python, and the Bartender1.1 barcode clustering algorithm^29^. Briefly, the raw sequences were kept for further analysis if they had the appropriate sequences for the sample index tags and the bases flanking the barcode regions, and if the mean quality score for the barcode sequence region was greater than 30. The Bartender1.1 clustering algorithm was then used to identify and count the barcodes from each sample. Barcode sequencing reads were then sorted based on the sample multiplexing tags and barcode read counts were corrected for PCR jackpotting effects^30^.

### Long-read sequencing

PacBio circular consensus HiFi sequencing was used to sequence the full Library Plasmid associated with each sensor. To achieve a high depth of long-read sequencing coverage of the full plasmid, two separate samples of the library plasmid DNA were prepared and sequenced. For both samples, the library plasmid DNA was extracted using miniprep kits. The plasmid DNA was linearized and dephosphorylated before submitting for sequencing (University of Maryland Institute for Genome Sciences).

Data was obtained from two PacBio Sequel II sequencing runs, with a total of 2 509 064 HiFi reads. The HiFi sequencing data was used to identify the *lacI* CDS for each sensor in the library by matching the *lacI* CDS to the corresponding DNA barcode. In addition, the full plasmid sequence was used to screen for sensors encoded in plasmids with unintended mutations in other plasmid regions that affected the measurement, and those sensors were excluded from quantitative phenotypic analyses.

### Fitness measurement

The experimental approach for this work was designed to maintain bacterial cultures in exponential growth phase for the full duration of the measurements. So, in all analysis, the Malthusian definition of fitness was used, i.e. fitness is the exponential growth rate^31^.

The fitness associated with each sensor was calculated from the change in the relative abundance of DNA barcodes over time. A spike-in control with known fitness was used to normalize the DNA barcode count data to enable the determination of the absolute fitness for each sensor in the library. The spike-in control was an *E. coli* clone containing a plasmid with a sensor with constitutively high *G*(*L*), i.e. the *I*^−^ phenotype^18^. The fitness of the spike-in control was determined from OD_600_ data acquired during growth of clonal cultures grown with the same automated growth protocol as used for the library fitness measurement. The fitness of the spike-in control was measured in all 24 growth environments and was independent of IPTG concentration but was slightly lower with tetracycline than without tetracycline. For each sensor in each of the 24 chemical environments, the ratio of the barcode read count to the spike-in read count was fit to a function assuming exponential growth and a lag in the onset of the fitness impact of tetracycline. The fitness associated with each sensor in each of the 24 chemical environments was determined as a parameter in the corresponding least-squares fit.

### Dose-response curve measurement

The Library Plasmid and Verification Plasmid were engineered to provide two independent measurements of the dose-response curve for LacI sensors. First, in the Library Plasmid, the sensor regulates the expression of a tetracycline resistance gene (*tetA*) that enables determination of the dose-response from barcode sequencing data by comparing the fitness measured with tetracycline to the fitness measured without tetracycline. Second, in the Verification Plasmid, the sensor regulates the expression of a fluorescent protein (YFP) that enables direct measurement of the sensor dose-response curve with flow cytometry.

A set of nine randomly selected sensors were used to calibrate the estimation of sensor output from the barcode-sequencing fitness measurements (Extended Data Fig. 9). The calibration data consisted of the fitness data for each calibration sensor from the library barcode sequencing measurement (using the Library Plasmid) and flow cytometry data for each calibration sensor prepared as a clonal culture (using the Verification Plasmid). This data was fit to a Hill equation model for the fitness impact of tetracycline as a function of the sensor output gene expression level, *G*:

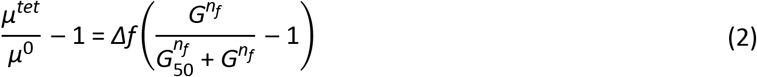

where *μ^tet^* is the fitness with tetracycline, *μ*^0^ is the fitness without tetracycline, *Δf* is the maximal fitness impact of tetracycline (when *G* = 0), *G*_50_ is the sensor output level that produces a 50 % recovery in fitness, and *n*_*f*_ characterizes the steepness of the fitness calibration curve. Because the fitness calibration curve, equation (2), is nonlinear, it cannot be directly inverted to give a sensor output value for all possible fitness measurements. So, two Bayesian inference models were used to estimate the dose-response curves for every sensor in the library using the barcode sequencing fitness measurements. Both inference models used equation (2) to represent the relationship between fitness and sensor output. The parameters *Δf*, *G*_50_, and *n*_*f*_ were included in both inference models as parameters with informative priors. Priors for *G*_50_ and *n*_*f*_ were based on the results of the fit to the fitness calibration data (Extended Data Fig. 9): *G*_50_ ~ normal(mean=13 330, std=500), *n*_*f*_ ~ normal(mean=3.24, std=0.29). We chose the prior for *Δf* based on an examination of μ^*tet*^⁄μ^0^ − 1 measured with zero IPTG: *Δf* ~ exponentially-modified-normal(mean=0.720, std=0.015, rate=14). The use of a prior for *Δf* with a broad right-side tail was important to accommodate sensors in the library for which μ^*tet*^⁄μ^0^ − 1 was systematically less than −0.722.

The first Bayesian inference model assumed that the dose-response curve for each sensor was described by the Hill equation (see equation (1), above). The Hill equation parameters for each sensor, *G*_∞_, *G*_0_, *EC*_50_, and *n* and their associated uncertainties were determined using Bayesian parameter estimation by Markov Chain Monte Carlo (MCMC) sampling with PyStan^32^. Broad, flat priors were used for log_10_(*G*_0_), log_10_(*G*_∞_), and log_10_(*EC*_50_), with error function boundaries to constrain those parameter estimates to within the measurable range (100 MEF ≤ *G*_0_, *G*_∞_ ≤ 50 000 MEF; 0.1 μmol/L ≤ *EC*_50,_*i* ≤ 40 000 μmol/L). For *ni*, we used a gamma distribution prior with shape parameter of 4.0 and inverse scale parameter of 3.33.

The second Bayesian inference model was a non-parametric Gaussian process (GP) model that assumed only that the dose-response curve for each sensor was a smooth function of IPTG concentration^33^. The GP model was used to determine which sensors had band-pass or band-stop phenotypes. The GP model was also implemented using MCMC sampling with PyStan^32^.

Source code for both models is included in the software archive given at the end of this manuscript. MCMC sampling for both models was run with 4 independent chains, 1000 iterations per chain (500 warmup iterations), and the adapt_delta parameter set to 0.9.

For quantitative analyses of sensor phenotypes based on Hill equation parameters, data were only included if the results of the Hill equation model and the GP model agreed. More specifically, data were only used for those analyses if the median estimate for the dose-response curve from the Hill equation model was within the central 90 % credible interval from the GP model at all 12 IPTG concentrations.

### Dose-response curve verification

Approximately 120 sensors from the library were chosen for flow cytometry verification of the dose-response curves. The CDSs of these sensors were chemically synthesized (Twist Bioscience), cloned into the Verification Plasmid, and then transformed into MG1655Δ*lac*. Transformants were plated in LB supplemented with kanamycin and 0.2 % glucose. Sensor sequences were verified with Sanger sequencing (Psomagen USA). For flow cytometry measurements of dose-response curves, a culture of *E. coli* containing the Verification Plasmid with a chosen sensor sequence was distributed across 12 wells of a 96-well plate and grown to stationary phase using the automated microbial growth system. After growth to stationary phase, cultures were diluted 50-fold into a plate containing the same 12 IPTG concentrations used during the landscape measurement (0 μmol/L to 2048 μmol/L). In some cases, higher IPTG concentrations were used to capture the full dose-response curves of selected sensors (e.g. Fig. 2i-j). Cultures were then grown for 160 minutes (~3.3 generations) before being diluted 10-fold into the same IPTG gradient and grown for another 160 minutes. Then, 5 μL of each culture was diluted into 195 μL of PBS supplemented with 170 μg/mL chloramphenicol and incubated at room temperature for 30-60 minutes to halt the translation of YFP and allow extant YFP to mature in the cells.

Samples were measured on an Attune NxT flow cytometry with autosampler using a 488 nm excitation laser and a 530 nm ± 15 nm bandpass emission filter. Blank samples were measured with each batch of cell measurements, and an automated gating algorithm was used to discriminate cell events from non-cell events. With the Attune cytometer, the area and height parameters for each detection channel are calibrated to give the same value for singlet events. So, to identify singlet cell events and exclude multiplet cell events, we applied a second automated gating algorithm that selected only cells with side scatter area ≅ side scatter height. All subsequent analysis was performed using the singlet cell event data. Fluorescence data was calibrated to molecules of equivalent fluorophore (MEF) using fluorescent calibration beads (Spherotech, part no. RCP-30-20A). The cytometer was programmed to measure a 25 μL portion of each cell sample, and the 40-fold dilution used in the cytometry sample preparation resulted in approximately 20 000 singlet cell measurements per sample. The geometric mean of the YFP fluorescence was used as a summary statistic to represent the output level of each sensor as a function of the input ligand concentration, [IPTG].

### Calculation of abundance for sensor phenotypes

The relative abundance of each sensor phenotype was estimated using the results of both Bayesian inference models (Hill equation and GP, Fig. 2a-b). Sensors were labeled as “flat response” if the Hill equation model and the GP model agreed (see above) and if the posterior probability for (*G*_0_ > *G*_∞_) was between 0.05 and 0.95 (from the Hill equation model inference). Sensors were labeled as having a negative response if the slope, ∂*G*/∂*L*, was negative at one or more IPTG concentrations with 0.95 or higher posterior probability (from the GP model inference). To avoid false positives from end effects, this negative slope criteria was only applied for IPTG concentrations between 2 μmol/L IPTG and 1024 μmol/L IPTG. Sensors were labeled as “always on” (the *I^−^* phenotype from reference^18^) if they were flat-response and if the sensor output level at zero IPTG was greater than 0.25 × *G*_∞,wt_ with 0.95 or higher posterior probability (from the GP model inference). Sensors were labeled as “always off” (the *I*^*S*^ phenotype from reference^18^) if they were flat-response but not always on. Sensors were labeled as band-stop or band-pass if the slope, ∂*G*/∂*L*, was negative at some IPTG concentrations and positive at other IPTG concentrations, both with 0.95 or higher posterior probability (from the GP model inference). Band-stop and band-pass sensors were distinguished by the ordering of the negative-slope and positive-slope portions of the dose-response curves. Sensors that had a negative response (see above) but that were not band-pass or band-stop, were labeled as inverted. False-positive rates were estimated for each phenotypic category by manually examining the fitness data for sensors with less than three substitutions. Typical causes of false-positive phenotypic labeling included unusually high noise in the fitness measurement and biased fit results due to outlier fitness data points. Estimated false-positive rates ranged between 0.001 and 0.005. The relative abundance values shown in Fig. 2a-b were corrected for false positives using the estimated rates.

### Comparison of synonymous mutations

The library contained a set of 39 sensors with the wild-type *lacI* CDS (but different DNA barcodes), and a set of 310 sensors with only synonymous nucleotide changes (*i*.*e*. no amino acid substitutions). Both sets had long-read sequencing coverage for the entire plasmid and were screened to retain only sensors with zero unintended mutations in the plasmid (i.e. no mutations in regions of the plasmid other than the *lacI* CDS). The Hill equation fit results for those two sets were compared to determine whether synonymous SNPs affected the measurable phenotype of a sensor. The Kolmogorov-Smirnov test was used to compare the distributions of Hill equation parameters between these two sets. The resulting p-values, 0.71, 0.40, 0.28, and 0.17 for *G*_0_, *G*_∞_, *EC*_50_, and *n* respectively, indicate that there were no significant differences between them. Additionally, the library contained 40 sets of sensors, each with four or more synonymous CDSs (including the set of synonymous wild-type sequences and 39 non-wild-type sequences). A hierarchical model was used to compare the Hill equation parameters within each set of synonymous CDSs. Within each set, the uncertainty associated with individual sensors was typically larger than the sensor-to-sensor variability estimated by the hierarchical model. Overall, these results indicate that synonymous SNPs did not measurably impact the phenotype of the sensor, so only the amino acid sequences were considered for any subsequent quantitative genotype-to-phenotype analysis.

### Analysis of single-substitution data

The single amino acid substitution results presented in Fig. 4 are a combination of direct experimental observations, DNN model results, and estimates of *G*_0_ for missing substitutions.

For direct experimental observations, multiple sensor variants were often present in the library with the same single substitution. For each SNP-accessible substitution, if there was only one sensor variant in the library, the median and standard deviation for each parameter were used directly from the Bayesian inference using the Hill equation model. If there was more than one sensor variant with a given single substitution, the consensus Hill equation parameter values and standard deviations for that substitution were calculated using a hierarchical model based on the eight schools model^34,35^. The hierarchical model was applied separately for each Hill equation parameter. The logarithm of the parameter values was used as input to the hierarchical model, and the input data were centered and normalized by 1.15 × the minimum measurement uncertainty. The standard normal distribution was used as a loosely informative prior for the consensus mean effect, and a half-normal prior (mean = 0.5, std = 1) was used for the normalized consensus standard deviation (i.e. hierarchical standard deviation). These priors and normalization were chosen so that the model gave intuitively reasonable results for the consensus of two sensor variants (i.e. close to the results for the sensor variant with the lowest measurement uncertainty). Results for the hierarchical model were determined using Bayesian parameter estimation by Markov Chain Monte Carlo (MCMC) sampling with PyStan^32^. MCMC sampling was run with 4 independent chains, 10 000 iterations per chain (5000 warmup iterations), and the adapt_delta parameter set to 0.975.

For *G*_0_, the direct experimental results were used for the 1047 substitutions plotted as gray points or red points and error bars in Fig. 4d. In addition, estimated values were used for the 83 missing substitutions that have been previously shown to result in an “always on” LacI phenotype (i.e., the *I*^*–*^ phenotype^18,19^). For these substitutions, plotted as pink-gray points and error bars in Fig. 4d, the median value was estimated to be equal to the wild-type value for *G*_∞_ (24 000 MEF), and the geometric standard deviation was estimated to be 4-fold, both based on information from previous publications^18,19^. Note that these 83 substitutions are completely missing from the experimental dataset, i.e. they are not found in any sensor variant, as single substitutions or in combination with other substitutions.

For *G*_∞_ and *EC*_50_, the direct experimental results were used for the 964 substitutions that are found as single substitutions in the library and that haver a consensus standard deviation for log_10_(*EC*_50_) less than 0.35. An additional 74 substitutions are found as single substitutions in the library, but with higher *EC*_50_ uncertainty. For these substitutions, either *EC*_50_ is comparable to or higher than the maximum ligand concentration used for the measurement (2048 μmol/L IPTG), or *G*_∞_ is comparable to *G*_0_ (or both). Consequently, the dose-response curve is flat or nearly flat across the range of concentrations used, and the Bayesian inference used to estimate the Hill equation parameters results in *EC*_50_ and *G*_∞_ estimates with large uncertainties. The DNN model can provide a better parameter estimate for these flat-response sensors because it uses data and relationships from the full library (e.g. the log-additivity of *EC*_50_) to predict parameter values for each single substitution. So, the DNN model results were used for these 74 substitutions. Finally, the DNN model results were used for an additional 953 substitutions that are found in the library, but only in combination with other substitutions (i.e. not as single substitutions).

### Identification of high-frequency substitutions and structural domains associated with inverted and band-stop sensors

The strongly inverted sensors discussed above and used for the plots in Extended Data Fig. 6a,c,e were identified by the following criteria: *G*_0_/*G*_∞_ ≥ 2, *G*_0_ > *G*_∞,*wt*_/2, *G*_∞_ < *G*_∞,*wt*_/2, and *EC*_50_ between 3 μmol/L and 1000 μmol/L. The strong band-stop sensors discussed above and used for the plots in Extended Data Fig. 6b,d,f were identified by the following criteria: *G*_0_ > *G*_∞,*wt*_/2, *G*_*min*_ < *G*_∞,*wt*_/2, and the slope, ∂log(*G*)/∂log(*L*), of less than −0.07 at low IPTG concentrations and greater than zero at higher IPTG concentrations, both with 0.95 or higher posterior probability (from the GP model inference). In addition, the sets of strongly inverted and strong band-stop sensors were manually screened for likely false positives due to outlier fitness data points.

A hypergeometric test was used to determine the substitutions that occur more frequently in the set of strongly inverted or strong band-stop sensors than in the full library. For each possible amino acid substitution, the cumulative hypergeometric distribution was used to calculate the probability of the observed number of substitutions in inverted or band-stop sensors under a null model of no association. This probability was used as a p-value for the null hypothesis that the number of inverted or band-stop sensors with that substitution resulted from an unbiased random selection of sensors. Substitutions were considered to occur at significantly higher frequency if they had a p-value less than 0.005 and if they occurred more than once in the set of inverted or band-stop sensors. In the set of strongly inverted sensors, ten associated (higher frequency) substitutions were identified: S70I, K84N, D88Y, V96E, A135T, V192A, G200S, Q248H, Y273H, and A343G. In the set of strong band-stop sensors, eight associated substitutions were identified: V4A, A92V, H179Q, R195H, G178D, G265D, D292G, and R351G. To estimate the number of false-positives, random sets of sensors were chosen with the same sample size as the strongly inverted (43) or the strong band-stop (31) sensors and the same significance criteria was applied. From 300 independent iterations of the random selection, the estimated mean number of false-positive substitutions was 2.1 and 2.3 for the inverted and band-stop sensors, respectively.

A similar procedure was used to determine which structural domains within the protein are mutated with higher frequency in the inverted or band-stop sensors. The structural domains considered included the secondary structure domains from the complete crystal structure of LacI^36^, as well as larger structural domains (N-terminal core, C-terminal core, DNA-binding, dimer interface) and functional domains (ligand-binding, core-pivot). The p-value threshold used for significance was 0.025. For the strongly inverted sensors, six domains were identified with a higher frequency of substitutions: the dimer interface, residues within 7 Å of the ligand-binding pocket, helix 5, helix 11, strand I, and the N-terminal core. For the strong band-stop sensors, three domains were identified: the C-terminal core, strand J, and helix 9. From 300 independent random selections of sensors, the estimated mean number of false-positive domains was 0.39 and 0.50 for the inverted and band-stop sensors, respectively.

### Deep neural network (DNN) modeling

The dataset was pruned to a set of high-quality sequences for DNN modeling. Specifically, data for a LacI sensor variant was only used for modeling if it satisfied the following criteria:

1. No mutations were found in the long-read sequencing results for the regions of the plasmid encoding kanamycin resistance, the origin of replication, the tetA and YFP genes, and the regulatory region containing the promoters and ribosomal binding sites for *lacI* and tetA (Extended Data Fig. 8).
2. The total number of barcode read counts for a sensor variant was greater than 3000.
3. The number of amino acid substitutions was less than 14.
4. The measurement uncertainty for log_10_(*G*_∞_) was less than 0.7.
5. The results of the Hill equation model and the GP model agreed at all 12 IPTG concentrations (see above).

After applying the quality criteria listed above, 47 462 sensor variants remained for DNN modeling.

Amino acid sequences were represented as one-hot encoded vectors of length L=2536, and with mutational paths represented as K × L tensors for a sequence with K substitutions. The logarithm of the Hill equation parameter values were normalized to a standard deviation of 1, and then shifted by the corresponding value of the wild-type sequence in order to correctly represent the prediction goal of the change in each parameter relative to the wild-type. A long-term, short-term recurrent neural network was selected for the underlying model^37^, with 16 hidden units, a single hidden layer, and hyperbolic tangent (tanh) non-linearities. Inference was performed in pytorch^38^ using the Adam optimizer^39^. For *EC*_50_ and *G*_0_, the contribution of individual data points to the regression loss were weighted inversely proportional to their experimental uncertainty. Model selection was performed with 10-fold cross-validation on the training set (80 % of all available data). Approximate Bayesian inference was performed with the Bayes-by-backprop approach^17^. Briefly, this substitutes the point-estimate parameters of the neural network with variational approximations to a Bayesian model, represented as a mean and variance of a normal random variable. Effectively, this only doubles the number of parameters in the model. A mixture of two normal distributions was used as a prior for each parameter weight, with the two mixture components having high and low variance respectively. This prior emulates a sparsifying spike-slab prior while remaining tractable for inference based on back-propagation. Posterior means of each weight were used to calculate posterior predictive means, while Monte-Carlo draws from the variational posterior were used to assess posterior predictive intervals.

## Data availability

The raw sequence data have been deposited in the NCBI Sequence Read Archive and are available under the project accession number PRJNA643436.

The processed data table containing information for each LacI variant in the library is publicly available via the NIST Science Data Portal, with the identifier ark:/88434/mds2-2259 (https://data.nist.gov/od/id/mds2-2259 or https://doi.org/10.18434/M32259).

## Code availability

All custom data analysis code is available at https://github.com/djross22/nist_lacI_landscape_analysis.

## Acknowledgements

We would like to thank Vanya Paralanov, Daniel V. Samarov, Aaron G. Kusne, and Swarnavo Sarkar for discussions during planning and execution of this work. We would also like to thank Jayan Rammohan, William B. O’Dell, Ben M. Scott, Zvi Kelman, and Elizabeth A. Strychalski for insights during the experimental work, as well as comments on the manuscript.

## Author contributions

D.S.T., and D.R. conceived of the process.

D.S.T, S.L., and D.R. developed the experimental workflow.

D.S.T. designed, built, and tested genetic constructs.

E.F.R., N.A., and D.R. developed and programmed automation protocols.

D.S.T., E.F.R., N.A., O.V., and D.R. performed landscape and verification experiments.

P.D.T. and D.R. performed Bayesian inference and model fitting.

P.D.T. designed and evaluated the recurrent architecture for machine learning.

P.D.T., N.D.O, and D.R. contributed to long-read sequencing analysis.

D.S.T., P.D.T, A.P., and D.R. wrote the manuscript.

All authors contributed to the manuscript.

## Disclaimer

The authors declare no competing interests

Certain commercial equipment, instruments, or materials are identified to adequately specify experimental procedures. Such identification neither implies recommendation nor endorsement by the National Institute of Standards and Technology nor that the equipment, instruments, or materials identified are necessarily the best for the purpose.

Correspondence and requests for materials should be addressed to David Ross.

## Notes

### Competing Interest Statement

The authors have declared no competing interest.

